# Genome size distributions in bacteria and archaea are strongly linked to phylogeny

**DOI:** 10.1101/2021.12.15.472816

**Authors:** Carolina A. Martinez-Gutierrez, Frank O. Aylward

## Abstract

The evolutionary forces that determine genome size in bacteria and archaea have been the subject of intense debate over the last few decades. Although the preferential loss of genes observed in prokaryotes is explained through the deletional bias, factors promoting and preventing the fixation of such gene losses remain unclear. Moreover, statistical analyses on this topic have typically been limited to a narrow diversity of bacteria and archaea without considering the potential bias introduced by the shared recent ancestry of many lineages. In this study, we used a phylogenetic generalized least-squares (PGLS) analysis to evaluate the effect of different factors on the genome size of a broad diversity of bacteria and archaea. We used dN/dS to estimate the strength of purifying selection, and 16S copy number as a proxy for ecological strategy, which have both been postulated to play a role in shaping genome size. After model fit, Pagel’s lambda indicated a strong phylogenetic signal in genome size, suggesting that the diversification of this trait is strongly influenced by shared evolutionary histories. As a predictor variable, dN/dS showed a poor predictability and non-significance when phylogeny was considered, consistent with the view that genome reduction can occur under either weak or strong purifying selection depending on the ecological context. Copies of 16S rRNA showed poor predictability but maintained significance when accounting for non-independence in residuals, suggesting that ecological strategy as approximated from 16S rRNA copies might play a minor role in genome size variation. Altogether, our results indicate that genome size is a complex trait that is not driven by any singular underlying evolutionary force, but rather depends on lineage- and niche-specific factors that will vary widely across bacteria and archaea.

**Author Summary:** The evolutionary forces driving genome size in bacteria and archaea have been subject to debate during the last decades. Independent comparative analyses have suggested that unique variables, such as the strength of selection, environmental complexity, and mutation rate, are the main drivers of this trait, which complicates generalizations across the Tree of Life. Here, we applied a phylogeny-based statistical approach to assess how tightly genome size is linked to evolutionary history in bacteria and archaea. Moreover, we also evaluated the predictability of genome size from the strength of purifying selection and ecological strategy on a broad diversity of bacteria and archaea genomes. Our approach indicates that genome size in prokaryotes is strongly dependent on phylogenetic history, and that genome size is the result of the interaction of variables like past events, current selection regimes, and environmental complexity that are clade dependent.

## Introduction

Bacterial and Archaeal genomes are densely packed with genes and contain relatively little non-coding DNA, and therefore an increase in genome size is directly translated into more genes [1–3]. In contrast, multicellular eukaryotes generally show genome expansion due to the proliferation of noncoding-DNA as a consequence of high genetic drift [2]. The absence of non-functional elements in prokaryotes is explained through the deletion bias process; newly acquired genes or existing genes are removed through deletions if selection on those genes is ineffective enough due to low selection coefficient [4–6]. Although narrowly constrained when compared with eukaryotes, prokaryotic genome sizes still vary by over an order of magnitude. Assuming an intrinsic deletion bias in prokaryotes, it remains unclear what evolutionary forces determine which genes are maintained and which are lost, and what determines the variability of genome sizes across the broad diversity of bacteria and archaea.

Multiple individual factors have been hypothesized to be primary drivers of genome size in bacteria and archaea. Early studies suggested that effective population size (Ne) may be the primary force that determines genome size in prokaryotes. For example, genome reduction has been observed in host-dependent microorganisms that have small Ne due to bottlenecks and therefore experience high levels of genetic drift. Under such evolutionary constraints, slightly deleterious deletions are accumulated and cause overall genome reduction [7–11]. Paradoxically, later studies focusing on abundant free-living planktonic lineages in the ocean suggested that genome reduction can also be observed in bacteria with large Ne that experience strong purifying selection [12–15]. Factors other than Ne and the strength of purifying selection have also been postulated to play a role in determining prokaryotic genome size. Recently, one study suggested that environmental stress leads to genome streamlining in soil bacteria [16], and that habitat complexity and ecological strategy therefore may also play a major role in determining genome size. The mutation rate has also been proposed to be a major factor that determines genome size [17,18]. In particular, it was suggested that a high mutation rate would cause the erosion of genes, loss of function, and subsequent reduction in genome size of streamlined and host-dependent microorganisms [17–19]. Given the large number of forces that have been proposed to be primary determinants of genome size, it remains largely unknown whether genome size in prokaryotes is driven by unique variables, their interaction, or variables that have specific influence depending on the lineage.

In order to explore the evolutionary forces driving genome size in bacteria and archaea, we tested several hypotheses using a collection of genomes encompassing a broad diversity of bacteria and archaea available on the Genome Taxonomy Database (Fig. 1) (Genome Taxonomy Database, GTDB, [20]). We first examined how strongly genome size is linked to prokaryotic phylogeny in order to evaluate whether the recent shared evolutionary history of many lineages may explain why some factors have previously been shown to be correlated with genome size. Because genome size has most commonly been viewed as a result of either effective population size or ecological niche [3,21], we also evaluated the use of dN/dS ratios and *rrn* operon copies as proxies, respectively. Lastly, we then examined the power of predictability of genome size from these variables using a phylogeny-based statistical approach that explicitly accounts for the evolutionary relationships between different taxa. Our work provides important insights into the complex mechanisms that shape genome size in bacteria and archaea.

**Figure 1.**
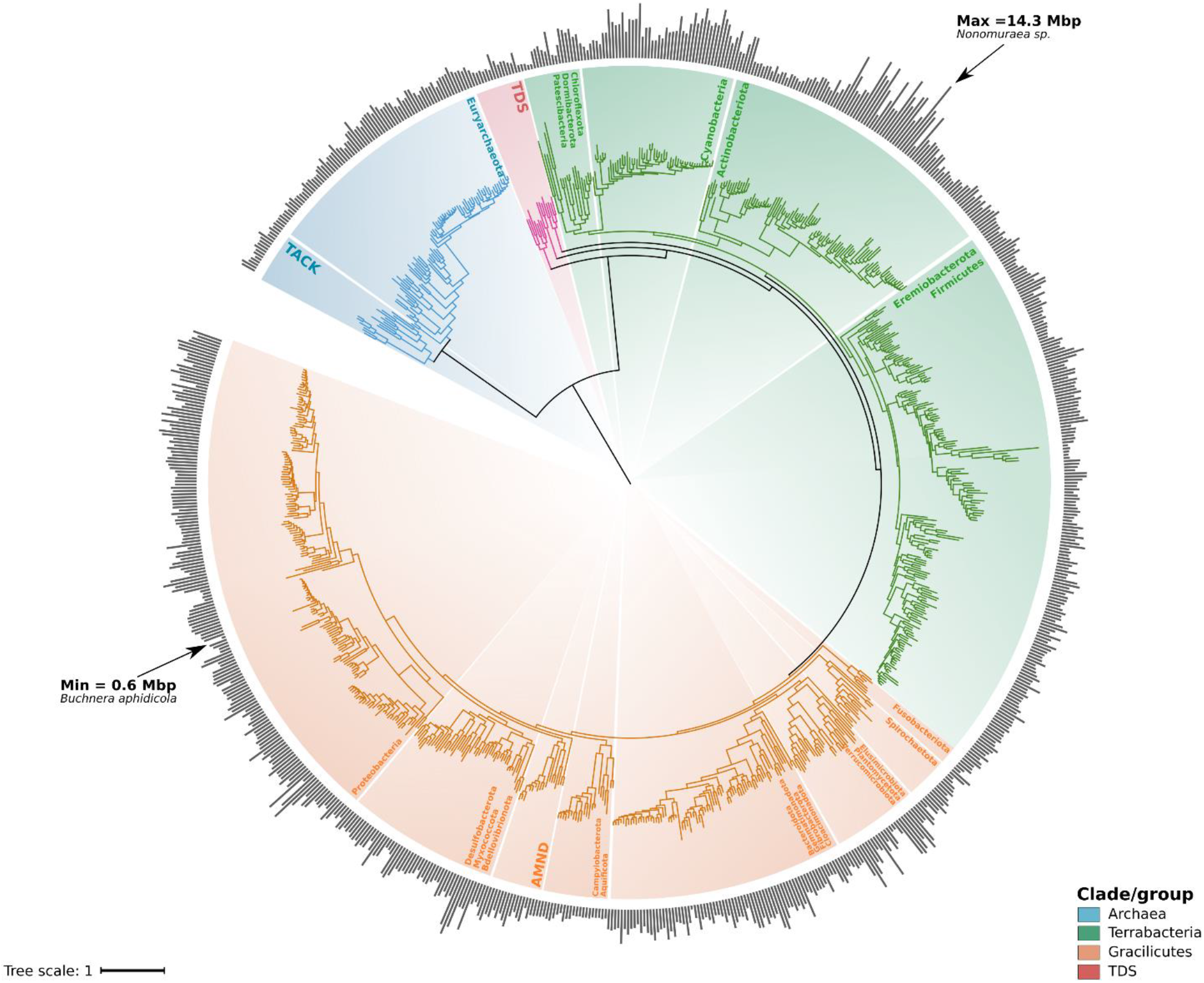
Genome size distribution across the Tree of Life of bacteria and archaea. Phylogenetic tree was built using a concatenated alignment of ribosomal and RNA polymerase sequences through a maximum likelihood approach and the substitution model LG+R10. Abbreviations: TACK = Thaumarchaeota, Aigarchaeota, Crenarchaeota and Korarchaeota; TDS = Thermotogota, Deinococcota, and Synergistota; AMND = Acidobacteriota, Methylomirabilota, Nitrospirota, Deferribacterota. Raw data for genome size can be found in Supplemental File S2.

## Results and discussion

### Genome size distribution across major phyla of bacteria and archaea

In order to explore the distribution of genome size across the Tree of Life of bacteria and archaea, we built a phylogenetic tree using representative genomes of 836 genera and 33 phyla available on the Genome Taxonomy Database [20] (Fig. 1, referred hereafter as GTDB genomes dataset. See methods for details on the criteria for genomes selection). For this phylogeny we used a set of ribosomal proteins and RNA polymerase subunits that we have previously benchmarked [22]. The size of genomes in our analysis and across the phylogeny varied by over one order of magnitude (0.6-14.3 Mbp, Fig. 1). The minimum and maximum corresponded to two bacterial lineages with contrasting lifestyles: the endosymbiont *Buchnera aphidicola* of the phylum Proteobacteria and the free-living Actinobacteria *Nonomuraea sp.* (Fig. 1). Regarding the distribution of genome size within each phylum, the greatest variation of genome size was observed for the phyla Actinobacteria and Cyanobacteria (Fig. 2A), whereas the phylum with shortest genomes belonged to symbiotic bacteria of the phylum Patescibacteria.

**Figure 2.**
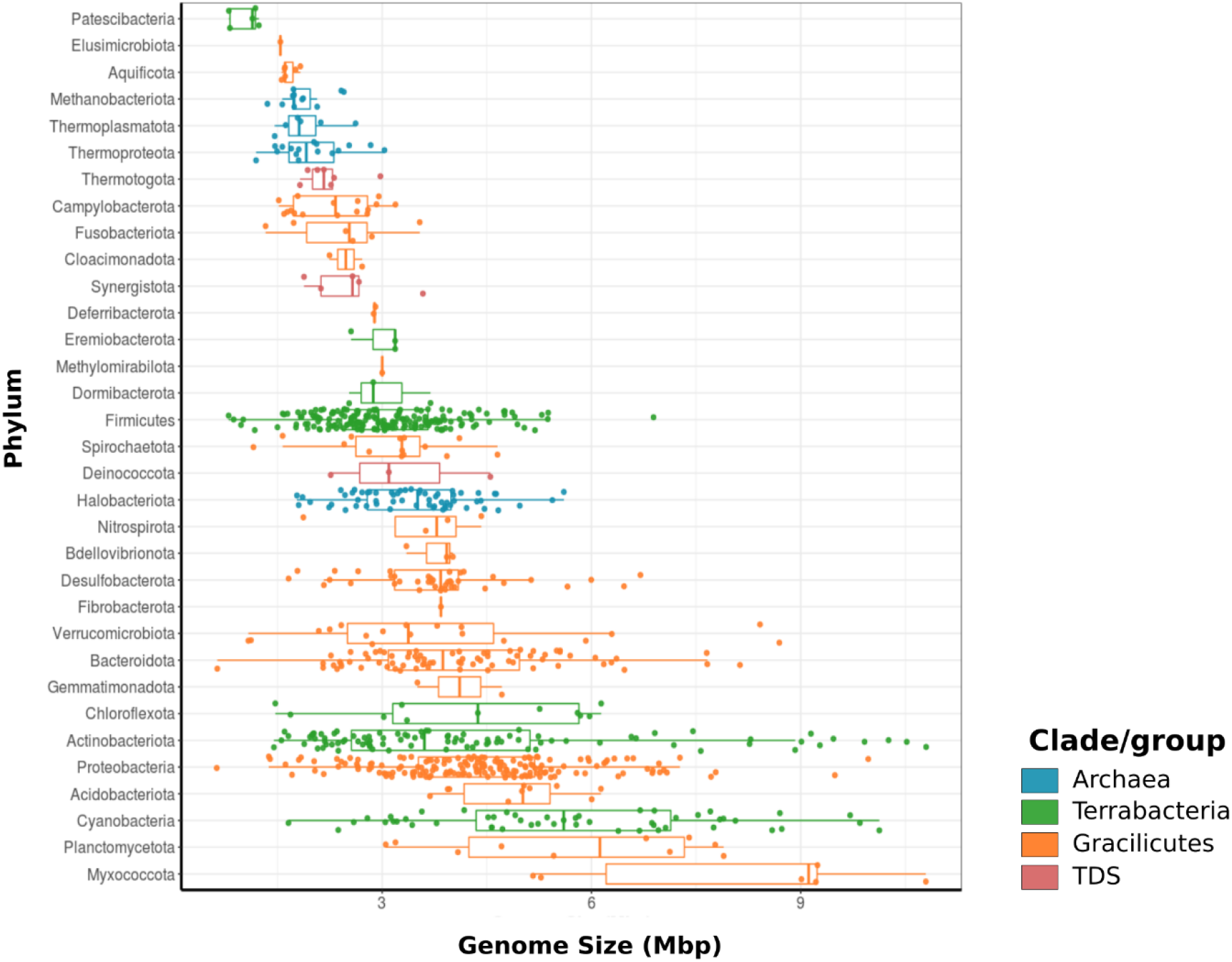
Distribution of genome size within bacteria and archaea taxonomic groups. Genome size grouping based on phylum. First, third quantile, and median are shown for each phylum distribution. Abbreviations: TDS = Thermotogota, Deinococcota, and Synergistota. Raw data for genome size can be found in Supplemental File S2.

Since the strength of selection and ecological strategy have been proposed to be important drivers of genome size in prokaryotes [21,23–25], we calculated the dN/dS and 16S rRNA copy number as approximations to the strength of selection and ecological strategy, respectively. We found that the largest genomes tended to have intermediate dN/dS values, while small genomes were found across a wide range of selection strengths (Fig. 3). These observations are consistent with previous descriptions of genome reduction occurring at high levels of purifying selection (i.e., *Pelagibacter* and *Prochlorococcus)* [15] but also under strong genetic drift (i.e., *Rickettsia, Blattabacterium,* and *Buchnera)* (Fig. 3) [21,26], indicating that there is not a strict linear relationship between genome size and selection strength. Similarly, we observed genomes with multiple 16S rRNA copies with variable dN/dS and genome size values (Fig. 3). Although we did not observe linear relationships between genome size and dN/dS or 16S rRNA copy number, we next sought to explore the predictability of this genomic trait from the latter parameters using a quantitative phylogenetic framework.

**Figure 3.**
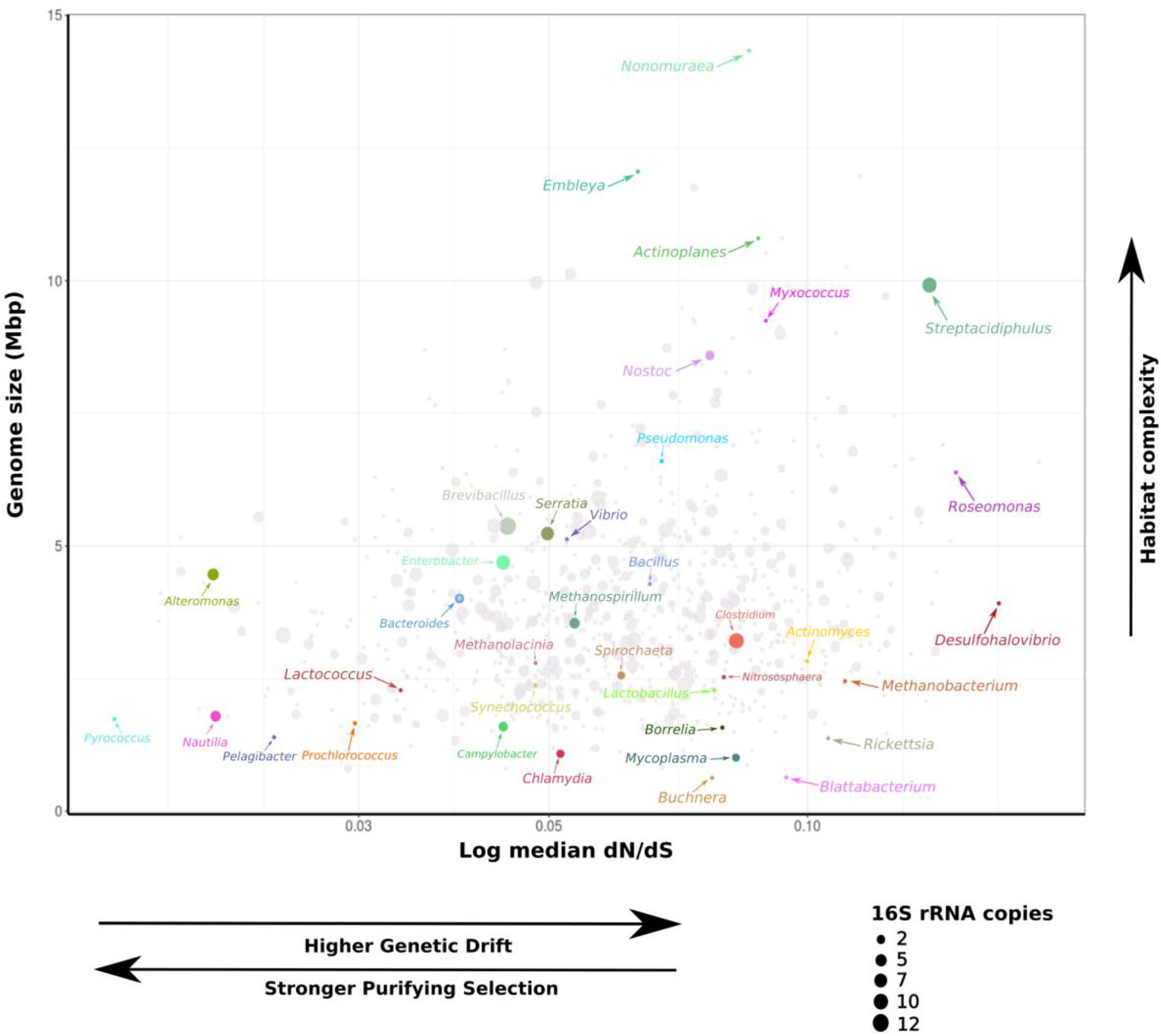
Relationship between genome size and dN/dS. dN/dS values were log transformed. Dots size is equivalent to the number of 16S rRNA gene copies. Raw data can be found in Supplemental File S2.

### Genome size is strongly dependent on phylogenetic history and clade-specific factors

Due to the recent shared evolutionary history of many bacteria and archaea, any study involving statistical analyses and species’ data potentially violates the assumption of independence of residuals [27,28], and phylogenetic methods are therefore required to analyse evolutionary relationships between traits. We sought to assess whether genome size distributions have a phylogenetic signal (i.e., that genome size is not randomly distributed across the Tree of Life and genome size variation is equivalent to phylogenetic distance). As a first approximation, we estimated Blomberg’s K [29] on the genome size of the GTDB genomes dataset (Figs. 1 and 2A). Values of Blomberg’s K between 0 and 1 indicate that the genome size of closely related genomes resemble each other, but less than expected under the Brownian Motion model (BM) of trait evolution, whereas K=1 is evidence that genomes size varies according to the Brownian Motion expectation [29]. We observed phylogenetic signal in the data, but less than expected under the Brownian Motion model (BM) (K=0.51, P=0.001), suggesting that although genome size in our data shows phylogenetic signal, variation is not fully explained through phylogenetic distance and relationships [30]. In addition, we tested the fit of different models of evolution for genome size, including Brownian Motion [30], Ornstein-Uhlenbeck [31], Early-Burst [32], a diffusion model, Pagel’s model [33], a drift model, and a white-noise model (non-phylogenetic signal) (Table 1). According to a likelihood ratio test (P<0.001 when compared with the next-best likelihood), the Pagel’s model showed the best fit (Table 1), supporting our previous finding that genome size in bacteria and archaea shows phylogenetic signal but it is not fully driven by phylogenetic history, which would be expected under the BM model.

**Table 1.** Summary of model fitting for genome size data. We highlighted the model that showed the highest likelihood and the lowest corrected AIC.

| Model | Loglik | AICc |
| --- | --- | --- |
| Brownian motion | -1463.3 | 2930.7 |
| Ornstein-Uhlenbeck | -1420.7 | 2847.4 |
| Early-Burst | -1463.3 | 2932.7 |
| **Pagel's lambda\*** | **-1415.6** | **2837.3** |
| Trend diffusion | -1447.7 | 2901.4 |
| Drift | -1463.3 | 2932.7 |
| White-noise | -1695.6 | 3395.2 |

\*Significantly higher likelihood when compared with the rest of the models tested according to the chisq test

We next used a phylogenetic generalized least squares model (PGLS) under Pagel’s approach to control for potential non-independence in the residuals of our regression models derived from the shared ancestry of the genomes analysed [34]. The Pagel’s lambda (λ) represents how strongly phylogenetic relationships predict the observed pattern of variation of a trait at the tips of a phylogeny and varies from 0 (no phylogenetic signal) to 1 (phylogenetic signal observed) [33]. According to our estimate, λ=0.98 (95% CI= 0.96-0.99; Table 2), genome size also shows non-phylogenetic independence in the residuals of the regression, confirming the suitability of a PGLS for the purposes of our analyses [35]. These findings indicate that phylogenetic history alone is a strong predictor of genome size, and that genome size in bacteria and archaea does not evolve independently. Similar results were obtained previously in a study analyzing the relationship between Ne*μ* and genome size but with a smaller set of prokaryotic and eukaryotic genomes [36,37], suggesting that sample size does not have an effect on the conclusions of our study. Moreover, we estimated kappa (k) and delta (δ), two parameters that describe the mode of evolution of a trait (punctuated vs gradual) and the rate change across the phylogeny (acceleration vs deceleration), respectively [38]. Our estimates (k=0.48-0.49 and δ=2.44-2.51, Table 2) are consistent with a gradual and late diversification of genome size in bacteria and archaea, which might indicate species-specific adaptations [33,38].

**Table 2.** Statistics of the regression models relating genome size and dN/dS and 16S rRNA as predictor variables using Generalized Least Square and Phylogenetic Least Square analyses. We highlighted the models that were statistically significant (α = 0.05).

| Model | Predictor variable | Kappa (95% CI) | Lambda (95% CI) | Delta (95% CI) | Slope | Intercept | P-val | AIC | R2* |
| --- | --- | --- | --- | --- | --- | --- | --- | --- | --- |
| Generalized Least Square |  |  |  |  |  |  |  |  |  |
| **Genome Size ~ Median dN/dS** | dN/dS | - | - | - | 13.57 | 2.97 | <0.001 | 3366.2 | 0.04 |
| **Genome Size ~ 16S rRNA copies** | 16S rRNA copies | - | - | - | 0.12 | 3.65 | 0.002 | 3387.7 | 0.01 |
| **Genome Size ~ Median dN/dS + 16S rRNA copies** | dN/dS + 16S rRNA copies | - | - | - | 14.11/0.13 | 2.7 | <0.001 | 3355.9 | 0.05 |
| Phylogenetic Generalized Least Square |  |  |  |  |  |  |  |  |  |
| Genome Size ~ Median dN/dS | dN/dS | 0.48<br>(0.39-0.58) | 0.98<br>(0.96-0.99) | 2.44<br>(2.01-2.85) | 1.26 | 2.46 | 0.5 | 2748.8 | 0.0006 |
| **Genome Size ~ 16S rRNA copies** | 16S rRNA copies | 0.49<br>(0.34-0.59) | 0.98<br>(0.96-0.99) | 2.49<br>(2.06-2.9) | 0.08 | 2.42 | 0.003 | 2740.268 | 0.01 |
| **Genome Size ~ Median dN/dS + 16S rRNA copies\*\*** | dN/dS + 16S rRNA copies | 0.49<br>(0.40-0.59) | 0.98<br>(0.96-0.99) | 2.51<br>(2.08-2.93) | 1.29/0.08 | 2.35 | 0.009 | 2741.79 | 0.01 |

\*R<sup>2</sup> calculated based on residuals, likelihood, and predicted data (multiple R-squared)
\*\*Anova did not show significant differences between models Genome Size ~ 16S rRNA copies and Genome Size ~ Median dN/dS + 16S rRNA copies (P=0.48)

### Non-phylogenetic regression overestimates the effect of dN/dS on genome size

Previous studies have suggested that high levels of genetic drift are related to a decrease in genome size in bacteria [21,26]. However, such studies were based on a limited set of genomes available at the time and did not include a broad repertoire of streamlined genomes, which are notable for their small genomes and large effective population sizes [10,39]. In order to investigate whether this trend is maintained when including a broader diversity of taxa, we calculated the dN/dS on the GTDB genomes dataset. Although earlier studies have reported a strong relationship between genome content and dN/dS [21], our non-phylogenetic generalized least squares (GLS) showed a positive and significant but poor predictability of dN/dS when using a broader set of genomes (P<0.001, R2=0.04, Table 1, Fig. 4A). Interestingly, when considering phylogeny, PGLS (phylogenetic generalized least squares) showed a non-significant and considerably poorer predictability (P=0.5, R2= 0.0006, Table 1, Fig. 4A). We also calculated the lambda parameter on dN/dS, and the value found (λ=0.68; 95% CI= 0.56-0.77) indicates a relatively high phylogenetic signal for this variable, suggesting that phylogenetically related microorganisms tend to experience similar levels of selection. Altogether, these results suggest that correlations between dN/dS and genome size are largely driven by artefacts that arise by not specifically accounting for the recent shared evolutionary history of many lineages [28].

**Figure 4.**
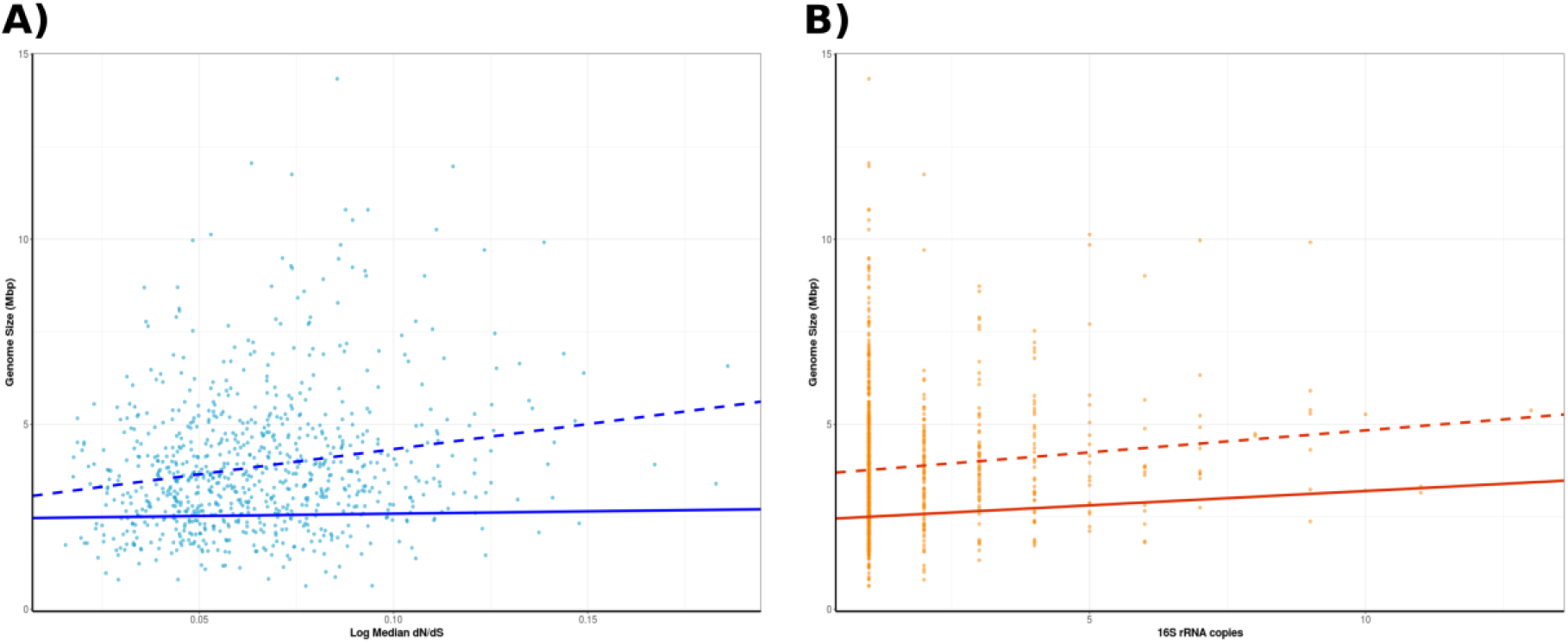
Relationship between genome size and genomic traits for bacteria and archaea. A) Regression line of the relationship between genome size and dN/dS ratio before (dashed line) and after (solid line) taking phylogenetic relationships into account. B) Regression line of the relationship between genome size and 16S rRNA copies before (dashed line) and after (solid line) taking phylogenetic relationships into account. Parameters of the regression equation for both relationships can be found in Table 2. Raw data can be found in Supplemental File S2.

Although our results indicate that dN/dS is a poor predictor of genome size in bacteria and archaea (Fig. 4A), it is worth mentioning that dN/dS only reflects recent evolutionary constraints due to saturation of substitutions at synonymous sites [40,41]. Therefore, we do not discount the possibility that genome reduction may be driven in part by processes such as population bottlenecks and periods of relaxed selection that happened in the past but are not reflected in dN/dS estimations. This scenario has been suggested for the streamlined autotroph *Prochlorococcus,* in which the genome simplification observed in this clade could be the result of periods of relaxed selection experienced in the past [41].

### Ecological strategy plays a role on genome size

In addition to testing the effect of strength of selection on genome size, we also assessed the predictability of genome size from 16S rRNA copies as an approximation to ecological strategy. Previous studies have shown that copies of the *rrn* operon can be a predictor of the number of ribosomes that a cell can produce simultaneously, and that this reflects the ecological strategy in microorganisms [25,42]. A large number of *rrn* copies is associated with the ability to adapt quickly to fluctuating environmental conditions (i.e., “boom and bust” strategies) [43], while multiple *rrn* copies would confer a metabolic burden to slow-growing microorganisms living in stable or low-nutrients environments because of ribosome overproduction [25]. Similarly to what we observed for dN/dS, we found a weak, positive, and significant relationship between genome size and 16S rRNA copies when using GLS (P<0.001, R2= 0.01, Table 2, Fig. 4B). However, our PGLS analysis did not reduce the R2 estimate when compared with the non-phylogenetic linear model (P=0.009, R2=0.01, Table 2, Fig. 4B), probably because the phylogenetic signal of 16S rRNA was relatively low (λ=0.40; 95% CI= 0.22-0.57). Although 16S rRNA copies show a poor predictability, our result suggests that larger genomes harbor more copies of the *rrn* open (Fig. 4B), consistent with the observation that larger genomes tend to inhabit complex environments in terms of temporal variability and diversity of resources [25]. In addition to fitting our model using dN/dS and 16S rRNA copies individually as predictors, we fitted an additive model with both variables. An ANOVA test showed that a model including both variables does not significantly improve the fit when compared with the model based on 16S rRNA copies as a unique predictor variable (P = 0.48).

### A hypothesis for the evolutionary processes that shape genome size in bacteria and archaea

According to our PGLS analysis (Table 1–2, Fig. 4), evolutionary history is a dominant variable determining genome size in bacteria and archaea, meaning that genomes with recent shared evolutionary history tend to maintain similar sizes since their divergence from their common ancestor. Nevertheless the pattern of variation in genome size differs from what would be expected under the Brownian Motion model, and microorganisms of the same clade can show variable genome size (Fig. 1–3). Based on our results we propose that genome size in prokaryotes is the result of a complex interplay of multiple variables, including evolutionary history, past events such as population bottlenecks, and environmental complexity (substrates available, variability in environmental factors, biotic pressure, etc.). The strong dependence of genome size on phylogenetic history suggests that when a group diverges, the resulting clades deviate from the genome size of the ancestor as a result of the colonization of new habitats, niche-specific adaptations, and/or population processes like bottlenecks or long population stability. Although several factors have been proposed to be singular drivers of genome size in prokaryotes, such as effective population size [44], ecological strategy [23], and mutation rate [17–19], our findings strongly suggest that genome size is a complex trait determined by lineage-specific factors that vary from group to group.

The phylogenetic signal detected in genome size does not discount that current and past processes like bottlenecks have a relevant role in the genome reduction of some bacteria and archaea. This is particularly expected in endosymbionts like *Buchnera* and *Blattabacterium,* which are thought to derive from a large-genome ancestor [8], and are frequently going through bottlenecks and periods of diversity loss [7,8,45]. Such exacerbated loss of diversity is enhanced by the nearly absent homologous recombination found in vertically transmitted endosymbionts [46]. These observations are consistent with the relatively high dN/dS value and small genome size that we observed for *Buchnera* and *Blattabacterium* (Fig. 3). In contrast, some abundant marine clades inhabiting the open ocean such as *Prochloroccocus* and *Pelagibacter* have undergone long periods of adaptation and specialization to their stable environments [47,48]. The open ocean is characterized by chronically-oligotrophic nutrient conditions that are stable throughout the year [49], and genes that are under relaxed selection are therefore pseudogenized and lost [10]. The latter is supported by the unusual growth requirements and low number of transcriptional regulators found in *Pelagibacter*, which is expected to limit its response to changing environmental conditions [50,51]. Consistent with these observations, we observed low dN/dS values, small genome size, and fewer 16S rRNA for these streamlined bacteria (Fig. 3). The small genomes observed in both endosymbionts and free-living planktonic lineages are therefore likely the result of distinct evolutionary processes, as previously proposed [15].

In contrast to the genome simplification observed in host-dependent and streamlined prokaryotes, genome expansion is expected in free-living lineages that inhabit complex environments like soils or sediments, where microenvironments with strikingly different abiotic conditions can be found millimeters apart [52]. Although temporal diversity declines and sweeps for specific gene variants are likely to occur in soil prokaryotes due to rapidly changing environmental conditions [52,53], larger genomes may be positively selected in these environmental realms due to variable abiotic and biotic constraints. Indeed, a study exploring the genes enriched in larger genomes of soil prokaryotes found a larger proportion of genes involved in regulation and secondary metabolism, and were depleted in genes related with translation, replication, cell division, and nucleotides metabolism when compared with smaller genomes [23]. These environmental and genomic findings are consistent with the large genome sizes, high dN/dS, and multiple 16S rRNA copies estimated in our study for soil microorganisms of the genera *Streptacidiphilus*, *Actinomyces, Conexibacter, Actinoplanes,* and *Myxococcus* (Fig. 3), the latter showing complex fruiting body development [54]. It is interesting to note that the largest genomes analyzed in our study (>6 Mpb) tend to experience intermediate levels of purifying selection (dN/dS), suggesting that either extremely high or low purifying selection are not conducive to genomic expansion events.

## Conclusions

Despite the increase of genomes available on publicly available databases, the evolutionary processes and factors driving genome size and content in bacteria and archaea are continuously debated. Several studies have proposed ecological strategies, the strength of purifying selection, and mutation rate as prominent forces that determine prokaryotic genome size, but our study shows that these factors likely vary in importance depending on the lineage. Moreover, our statistical approach showed that evolutionary history plays a large role in structuring genome size distributions across the Tree of Life, and that genome size is not a phylogeny independent trait. The significant but poor relationship between genome size and 16S rRNA copies suggest that besides phylogenetic history, ecological strategy plays a role in shaping genome size in bacteria and archaea, although this single trait is insufficient to completely represent ecological strategies.

Future studies will be necessary to evaluate this in detail on a lineage-by-lineage basis. he strong phylogenetic signal observed in genome size data indicates that analyses involving this trait cannot consider species as phylogenetically independent, therefore phylogenetic relatedness should be taken into account when studying the evolutionary forces driving genome size in order to avoid biased association between traits and simplified models.

## Material and methods

### Genomes compilation, dN/dS estimation, and *rrn* genes identification

In order to assess the predictability of genome size (response variable) from dN/dS and 16S rRNA copies (predictor variables), all the bacteria and archaea representative genomes available on the Genome Taxonomy Database (GTDB) (Release 05-RS95; 17th July 2020) [55] were filtered based on completeness (>=95%) and contamination (<=5%) and then classified at the Class levels. In order to include the phylum *Patescibacteria* in our analysis (also known as Candidate Phyla Radiation or CPR), we used completeness>=80% and contamination<=5% for this taxa. Classes having more than 500 genomes were randomly downsample to 500 genomes. The resulting genomes were clustered based on their taxonomic identity at the genus level. Genera with fewer than two genomes after filtering and clustering were discarded from further analyses. To estimate the strength and direction of selection on the genomes analysed, we calculated the ratio of synonymous and nonsynonymous substitutions (dN/dS) within each genus cluster using two sets of conserved marker genes, checkm_bact and checkm_arch for bacteria and archaea, respectively [56]. Genomes used to calculate the dN/dS for each genus cluster are reported in Supplemental File 1. The open reading frames (ORFs) retrieved from the GTDB were compared to the HMMs of the checkm_bact (120 marker genes) and checkm_arch marker (122 marker genes) sets using the hmmsearch tool available in HMMER v. 3.2.1 with the reported model-specific cutoffs [57]. We aligned the amino acid sequences for each marker gene and each cluster individually using ClustalOmega [58] and then converted amino acid alignments into codon alignments using PAL2NAL with the parameter --nogap [59]. We used the resulting codon alignments to estimate the ratio of synonymous and nonsynonymous substitutions for each pair of genomes using the maximum likelihood approximation (codeML) available on PAML 4.9h [60]. In order to avoid bias associated with divergence, dN/dS estimates with dS>1.5 were removed due to potential saturation. We also discarded pairwise comparisons with dS<0.1 because these might represent dN/dS values calculated from genomes of the same population. Moreover, dN/dS values >10 were considered artifactual [39]. Genomes with fewer than 25 marker genes remaining after filtering were discarded. After dN/dS estimation, we randomly selected one representative genome for each genus for further analyses (GTDB genomes dataset). We predicted ribosomal RNA genes in our selected genomes using Barrnap (barrnap 0.9: rapid ribosomal RNA prediction; https://github.com/tseemann/barrnap), with the default parameters. Genome size, 16S rRNA copies, and dNdS values for the GTDB representative genomes dataset are reported in Supplemental File S2.

### Statistical analyses

Due to the tendency of related species to resemble each other because of their shared phylogenetic ancestry, we assessed the suitability of a phylogeny-based method for our regression analyses by first estimating Blomberg’s K on genome size data [29] using the phylosignal function on R [61]. This parameter represents the phylogenetic signal in a continuous trait, and goes from 0 (no phylogenetic signal) to ∞ (phylogenetic signal) with the null hypothesis (K=1) meaning that the trait analysed evolves under Brownian Motion (BM, variation of the trait is proportional to the distance between species [30,62]. In addition, we also tested the fit of different trait evolution models, including including Brownian Motion [30], Ornstein-Uhlenbeck [31], Early-Burst [32], a diffusion model, Pagel’s model [33], a drift model, and a white-noise model (non-phylogenetic). We also performed a Generalized Least Square analysis to explore the predictability of genome size using dN/dS and 16S rRNA copies as predictor variables using the “glm” function available on R. Since we detected phylogenetic signal in genome size data, we additionally accounted for potential phylogenetic nonindependence in the residuals using the PGLS method with the function pgls on the R package Caper [63] and the Pagel’s model [33]. We calculated the lambda (λ) parameter (which showed phylogenetic signal in the residuals), delta (δ) and kappa (κ) (pattern of evolution of trait) through maximum likelihood. The best fitting model according to AIC and likelihood was checked visually using diagnostic plots (residuals vs. fitted values, and QQ plots to check normality) (Fig. S3).

### Phylogenetic reconstruction

To perform a Phylogenetic Generalized Least Square analysis (PGLS), we reconstructed a phylogenetic tree using the GTDB genomes dataset described above. We used the MarkerFinder pipeline reported previously [22], consisting in the identification of 27 ribosomal proteins and three RNA polymerase genes [64] using HMMER3 and the resulting individual sequences aligned with ClustalOmega and concatenated. In addition, the concatenated alignment was trimmed with trimAl [65] using the option -gt 0.1. The Ribosomal-RNAP alignment was then used to build the phylogenetic tree with IQ-TREE 1.6.12 [66] with the substitutions model LG+R10 and the options -wbt, -bb 1000, and --runs 10 [67–69]. The resulting phylogeny was manually inspected on iTOL [70] (Fig. 1).

## Acknowledgments

We acknowledge the use of the Virginia Tech Advanced Research Computing Center for bioinformatic analyses performed in this study. This investigation was supported by grants from the Institute for Critical Technology and Applied Science and the National Science Foundation (IIBR-1918271), and a Simons Early Career Award in Marine Microbial Ecology and Evolution to F.O.A. We kindly thank members of the Aylward Lab for their insightful comments on an earlier version of this manuscript and Prof. Josef Uyeda for advice on phylogeny-based statistical methods.

---

**Supplementary File 1.** Genomes used to calculate pairwise dN/dS within each genus cluster.

**Supplementary File 2.** dN/dS, genome size, and *rrn* operon copies for each genus representative.

**Supplementary Figure 3.** Diagnostic plots for the PGLS model genome size ~ 16S rRNA copies.

## Notes

### Competing Interest Statement

The authors have declared no competing interest.

