## Supplementary figures and images for "Genome size distributions in bacteria and archaea are strongly linked to phylogeny"

### Supplemental File 3

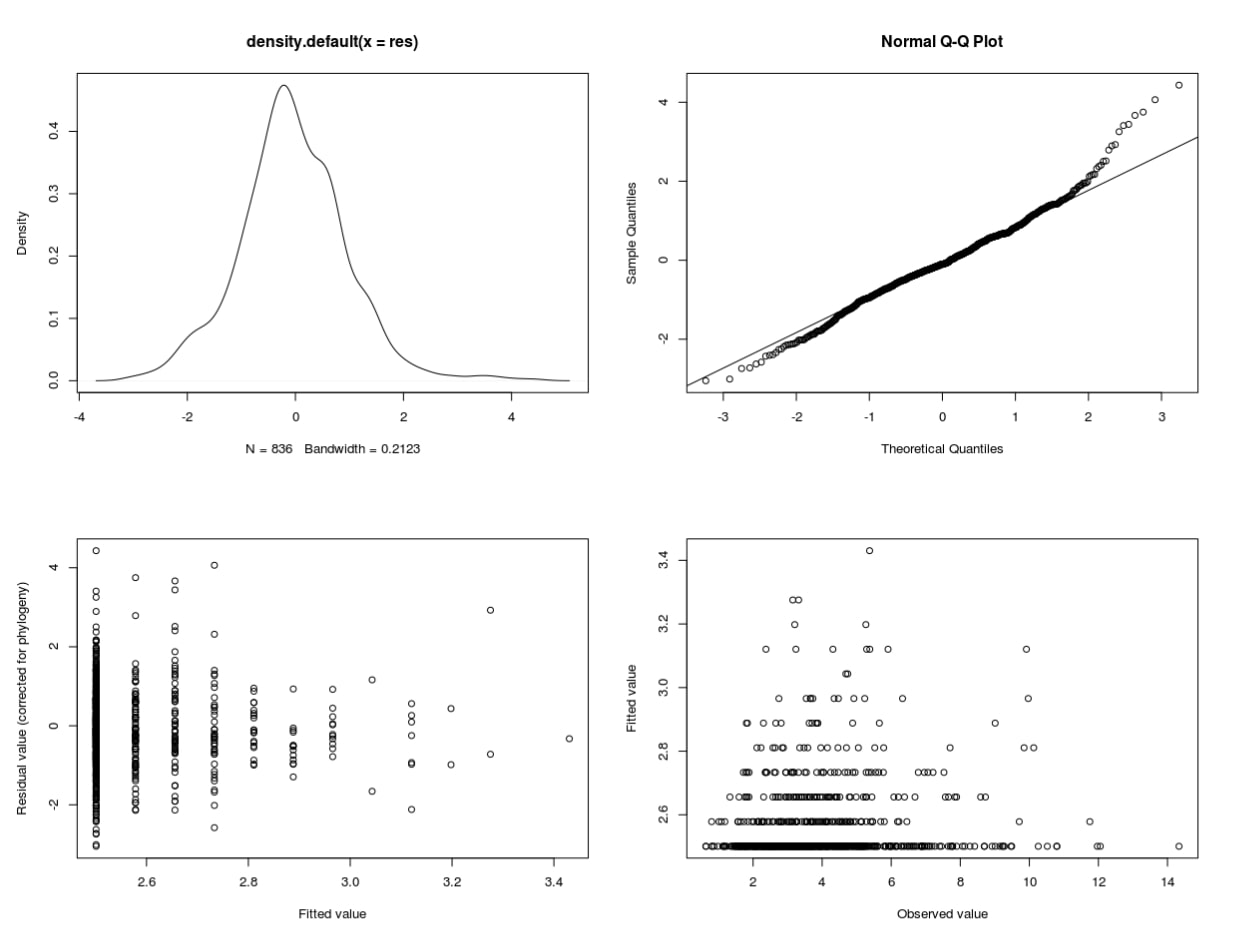
